# Wolf microevolution in the melting pot: range expansion, population sympatry, dynamic mosaic admixture zone and asymmetric gene flow

**DOI:** 10.64898/2026.09.02.748779

**Authors:** Jana Šrutová, Nikola Tkáčová, Ema Cetkovská, Kristýna Eliášová, Lenka Veselovská, Lenka Ungrová, Kamila Montoya, Michal Škrobánek, Petr Matějů, Martin Duľa, Aleš Vorel, Jan Mokrý, Karolína Mikslová, Sebastien Collet, Carsten Nowak, Francesca Rolle, Francesca Marucco, Maciej Szewczyk, Robert Mysłajek, Sabina Nowak, Slavomír Finďo, Vladimír Antal, Miroslav Kutal, Jindřiška Jelínková, Barbora Černá Bolfíková, Pavel Hulva

**Author notes:** **Corresponding author** Pavel Hulva.

## Abstract

Here we describe substantial range shifts of historically differentiated wolf populations and formation of novel sympatric zones during the recent decade in Central Europe. This region provides a natural laboratory for testing alternative scenarios of population interactions from continued isolation or restricted gene flow to progressive population fusion, while prompting a reassessment of their geographic ranges. Based on sampling across the Czech Republic and Slovakia obtained from large-scale monitoring programmes over five wolf years (2020/21–2024/25), complemented by comparative material from neighbouring regions, we analysed mitochondrial haplotypes, autosomal microsatellite genotypes and sex-linked loci. The Central European population with Baltic ancestry predominated across large parts of Central Europe including the Bohemian Massif with enclaves in the Western Carpathians. The Carpathian population was predominant in Slovakia, with a smaller satellite occurrence in the northern part of the Bohemian Massif. Alpine population was centred in the Alps but extended into southern parts of Bohemian Massif and Central German Uplands. Following the sporadic occurrence of admixed individuals, broad mosaic and dynamic sympatric zones have formed in the Czech Republic and Slovakia in the last decade. These scenarios could be facilitated by the presence of intermediate habitats and isolation of the Bohemian Massif structural basin, framed by a massive ring fault system. Recent-immigration estimates are asymmetric, with the largest mean contributions from the Alpine to the Central European population and from the Central European to the Carpathian population, with the second case potentially linked to source-sink dynamics driven by the hunting pressure within the Carpathian population (whereas the others are protected year-round). Whether increasing admixture will enhance viability of populations (that currently have small effective sizes) through genetic rescue or carry risks of outbreeding depression remains uncertain, highlighting the need for continued transboundary monitoring within conservation biology framework.

## Introduction

The recovery of large carnivores is one of the most prominent examples of wildlife restoration within the human-dominated landscapes of Europe. During the past centuries, large carnivores were severely depleted or extirpated across much of Europe, primarily as a consequence of direct persecution, habitat transformation and depletion of wild prey (Boitani, 2003; Ripple et al., 2014). The grey wolf (*Canis lupus*) was formerly widespread across the continent but disappeared from large parts of Western and Central Europe, persisting in fragmented populations, particularly on the Iberian and Apennine Peninsula, the Balkans, the Carpathians, the Dinaric region and Eastern Europe (Boitani, 2003; Hindrikson et al., 2017; Randi, 2011). Since the late twentieth century, legal protection and increasing societal tolerance have enabled wolves to recolonise many human-dominated landscapes, while farmland abandonment and forest regeneration have increased habitat and prey availability in several parts of Europe (Chapron et al., 2014; Navarro & Pereira, 2012). This trend stands in contrast to most other parts of the world, where large animals including large carnivores are generally in decline (Ingeman et al., 2022; Ripple et al., 2014). This makes Europe a global laboratory for research on megafauna recovery, which is of particular relevance within the context of rewilding, which emphasises the restoration of ecological interactions and landscape connectivity (Carver et al., 2021; Soulé & Noss, 1998).

### Genetic consequences of recolonization and range expansion

From a population genetic perspective, recolonization is not solely a demographic process. Serial founder effects, genetic drift and allele surfing may alter allele frequencies during range expansion, different selective regimes may occur in novel areas, whereas long-distance dispersal and secondary contacts may re-establish gene flow and generate admixed populations (Fabbri et al., 2007; Hulva et al., 2024; Szewczyk et al., 2019). Consequently, recently recolonised areas may contain complex genetic mosaics rather than representing simple geographic extensions of a single source population. European wolves retain pronounced mitochondrial and nuclear genetic structure resulting from historical phylogeographic processes, population fragmentation and recent demographic change (Hindrikson et al., 2017; Pilot et al., 2010; Stronen et al., 2013). Understanding how differentiated populations interact during contemporary range expansion is therefore important for identifying conservation units and assessing their future microevolutionary trajectories and viability. Three major wolf populations are particularly relevant to recolonization in Central Europe: the Central European, Carpathian and Alpine populations. These populations originate from refugia in distinct biomes, likely bear the signatures of Pleistocene phylogeographic differentiation (Pilot et al., 2014; Silva et al., 2020), and differ in mitochondrial haplotype composition and autosomal genetic structure (Czarnomska et al., 2013; Hindrikson et al., 2017; Stronen et al., 2013).

### Interpopulation admixture within Central Europe

The Central European population was founded from the Baltic population by a relatively small number of long-distance dispersers, and subsequently expanded westward through Poland and into adjacent regions of Germany (Jarausch et al., 2021; Szewczyk et al., 2019). Genetic analyses of Polish wolves identified differentiation between Central European and Carpathian populations in both mitochondrial and microsatellite data, with evidence of admixture and dispersal between them (Czarnomska et al., 2013). The Carpathian population has persisted in eastern Central Europe and remains an important and genetically differentiated component of the regional wolf population (Hindrikson et al., 2017; Pilot et al., 2010; Stronen et al., 2013). In parallel, the Alpine population predominantly derives from the natural expansion of the Italian Apennine population into the western Alps, a process accompanied by founder effects and initially restricted gene flow (Fabbri et al., 2007; Lucchini et al., 2002). Populations that extend into Central Europe only marginally, such as the Pontic and Dinaric-Balkan populations, are not analysed in detail in present study.

Contact between Central European and Carpathian wolves has been detected in the Western Carpathians and adjacent regions, supporting the interpretation of Central Europe as a phylogeographic interface characterised by dynamic range expansion and population contact (Hulva et al., 2018; Szewczyk et al., 2019), which fits into textbook phylogeographic paradigms regarding the role of this region as a suture zone for different lineages (Hewitt, 2000). More recently, admixture between Central European and Alpine wolves was documented, for instance, in the Bohemian–Bavarian Forest, demonstrating that expansion from the Alps has contributed to the genetic composition of wolves beyond the Alps (Hulva et al., 2024). Admixture may increase local genetic diversity and restore connectivity among formerly isolated populations, but it may also produce complex patterns of individual ancestry, admixture and contemporary gene flow that cannot be inferred from geographic origin alone.

### Study rationale and aims

Contact between previously isolated and differentiated populations can result in a range of scenarios, spanning from the maintenance of population isolation and “isolation with migration” to population fusion. Factors promoting fusion are linked to the fact that this involves intraspecific variability, whereas factors maintaining population distinctness include population differentiation and potential environmental / ecotypic adaptations, as well as phenomena such as natal habitat dispersal (Sanz-Pérez et al., 2018). Given the intraspecific divergence of the studied populations - which, however, occur in a mosaic of heterogeneous environments - we hypothesise that the initial stages of contact will involve intermediate scenarios and temporal steady state of the “isolation with migration” type.

In addition to previous qualitative studies, extensive monitoring programs conducted in collaboration with state and non-profit nature conservation organizations in the Czech Republic and Slovakia make it possible, for the first time, to obtain the quantitative data needed to test the aforementioned hypotheses. For this purpose, we utilised mitochondrial DNA data, autosomal microsatellite genotypes and sex markers and population genetics methods to investigate the population composition of recolonizing wolves in Central Europe. Specifically, we aimed to: (i) obtain information about population structure and distribution of particular clusters / populations; (ii) compare regional patterns of allelic richness and private allelic richness as complementary measures of genetic differentiation; (iii) identify secondary contact zones and proportion of individuals showing mixed ancestry; and (iv) evaluate individual assignment to candidate reference populations and estimate recent immigration among the principal wolf populations.

## Material and methods

### Study area, sampling and range mapping

The main portion of genetic material was collected during routine monitoring of grey wolves in the Czech Republic and Slovakia over five consecutive wolf years (WY; 1 May–30 April), from WY 2020/21 to 2024/25 (n = 3,818, Supplementary Data S1). Sampling involved governmental and non-governmental conservation organisations, research teams and trained citizen scientists, with collection procedures standardised through in-person and online training. Non-invasive samples included scat, hair, urine, blood deposited by females during oestrus and swabs from livestock carcasses. Tissue samples were obtained from wolf mortalities, mainly road-killed or illegally killed individuals. Samples were stored in 96% ethanol at −20 °C. Reference samples represented the Carpathian population in Slovakia and Poland (n = 323), the Central European population in Germany and Poland (n = 106), and the Alpine population (n = 28), mainly from Italy, as well as published mitochondrial records (Supplementary Data S1).

Sampling locations of focal and reference samples in relation to the distribution of European wolf populations are shown in Fig. S1, displayed within the EEA (European Environment Agency) regular square grid (10 × 10 km) according to latest published distributional data (Kaczensky et al., 2024), actualised by new records from present study. Maximum range extents of particular populations were displayed as extent of occurrence (EOO; IUCN Standards and Petitions Committee, 2024), represented by minimum convex polygons (MCPs; Fig. 1a). These MCPs were based on empirical data including permanent and sporadic occurrence records and were created using records of each population’s most frequent mtDNA haplotypes: HW01 and HW02 for the Central European population, HW06 and HW14 for the Carpathian population, and HW22 for the Alpine population, categorised sensu Pilot et al., 2010. Rare haplotypes were not interpreted. More detailed mapping of contact zones within the EEA grid was performed using an orthogonal convex hull based on the distribution of foreign mtDNA haplotypes (representing dispersing animals or backcrosses with mtDNA introgression) or admixed nuclear genotypes with q > 0.75 (corresponding to dispersing animals or first backcross generation). For population genetics approaches requiring a priori defined groups, multilocus genotypes were assigned to six operational geographic sampling regions based on wolf habitat environmental clusters (Hulva et al., 2018) and their potential geographic isolation: the Alps, Central German Uplands, Central European lowland, Baltic region, Bohemian Massif and the Carpathians (Fig. 1b).

**Fig. 1.**
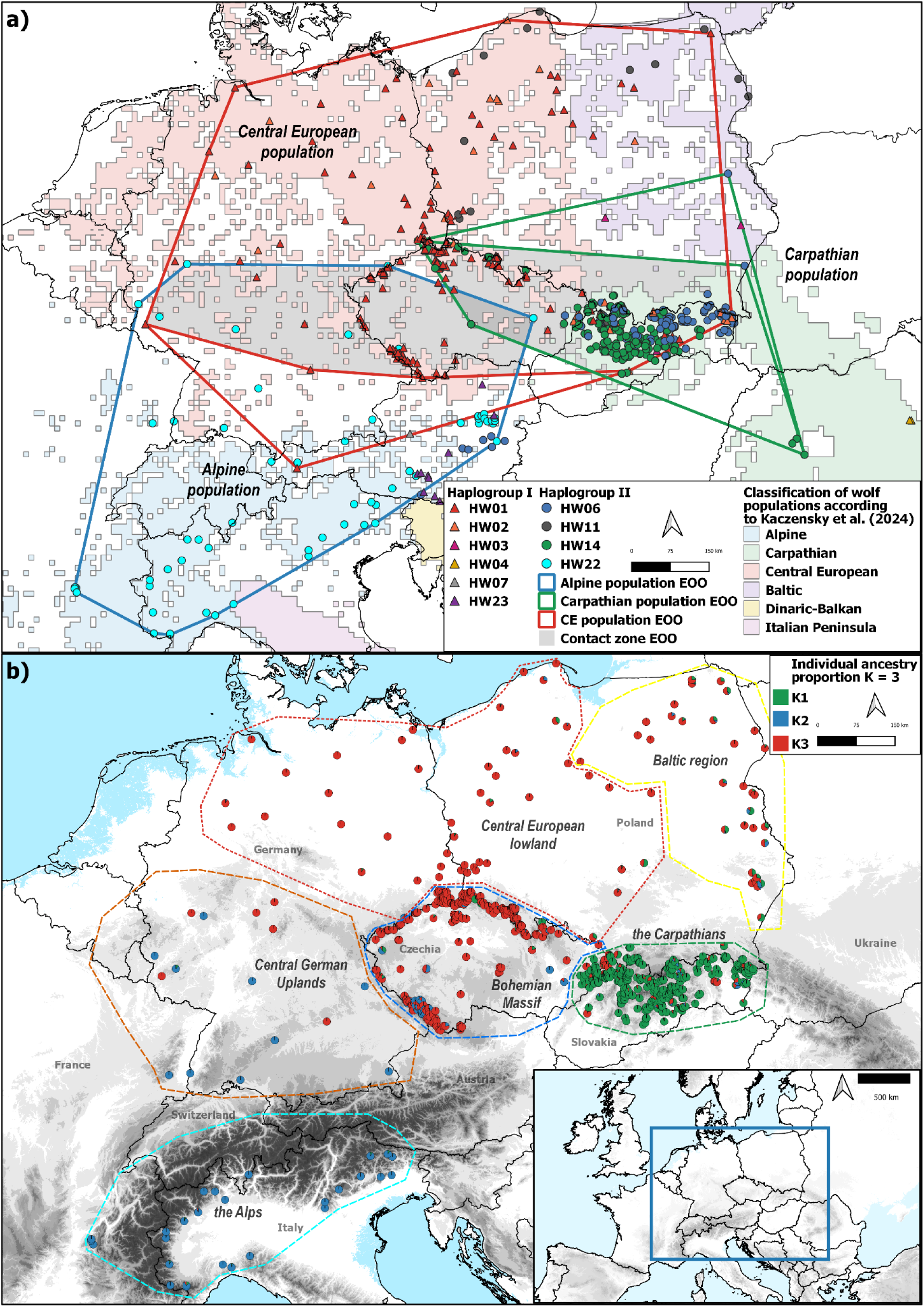
Ranges of studied grey wolf populations and spatial distribution of mitochondrial and nuclear data. (**a**) Individual mtDNA haplotypes are shown by coloured symbols. Maximum range extents of particular populations were displayed as extent of occurrence (EOO), represented by minimum convex polygons (MCPs) created by merging data from records of population-typical mtDNA haplotypes with triangles representing haplogroup 1 and circles haplogroup 2. Population characterised sensu Chapron et al., 2014 and their current distribution displayed in EEA square grid sensu Kaczensky et al., 2024. (**b**) Spatial distribution of individual Structure membership coefficients for K = 3 in the relatedness-pruned dataset. Each pie chart represents an individual, with sector proportions showing ancestry coefficients inferred from multilocus microsatellite genotypes. Colours denote membership in the predominantly Carpathian, Alpine and Central European genetic component. Dashed polygons delimit the six predefined geographic sampling regions. Grey tones are scaled by 250 m of altitude.

### DNA extraction and mitochondrial DNA sequencing

DNA was extracted using sample-specific commercial kits: QIAamp^®^ Fast DNA Stool Mini or NucleoSpin^®^ DNA Stool kits for scat, QIAamp^®^ Fast DNA Stool Mini Kit for urine, DNeasy^®^ Blood and Tissue or Genomic DNA Mini Kit (Tissue) for tissue, hair and livestock-carcass swabs, and Blood/Cell DNA Mini Kit for oestrus-blood samples. Sample preparation included extraction from the outer layer of scat, ethanol precipitation of urine, digestion of tissue pieces up to 25 mg and use of hair fragments containing follicles. Detailed extraction conditions are provided in Supplementary Methods S1. Subsequent steps followed the manufacturers’ protocols, and DNA was eluted in 30–100 µl of the respective elution buffer.

Mitochondrial haplotypes for a subset of samples were determined from the left hypervariable domain of the control region, amplified following Hulva et al. (2018), Jarausch et al. (2021) and Hulva et al. (2024) using primers Thr-L15926 and DL-H16340 (Vilà et al., 1999). The resulting 230-bp fragment was used for haplotype identification (Pilot et al., 2010) and, where microsatellite data were insufficient or ambiguous, species verification. PCR, amplicon purification and electrophoresis conditions are provided in Supplementary Methods S1. Forward sequencing with Thr-L15926 was performed on a 3130xl Genetic Analyzer (Applied Biosystems), and chromatograms were inspected and edited in Geneious (Kearse et al., 2012).

### Microsatellite genotyping

Twenty autosomal microsatellite loci and the sex-determining locus Amelogenin were co-amplified in two multiplex PCRs using fluorescently labelled primers and the Multiplex PCR *Plus* Kit (QIAGEN) following Hulva et al. (2024). Each 10 µl reaction contained 5 µl of Multiplex PCR Plus Master Mix, 3 µl of nuclease-free water, 1 µl of multiplex primer mix, and 1 µl of template DNA. Marker characteristics are provided in Table S1. PCR cycling comprised an initial denaturation at 95 °C for 5 min, followed by 35 cycles of 94 °C for 30 s, 56 °C for 90 s and 72 °C for 60 s, and a final extension at 72 °C for 10 min. Products were separated by capillary electrophoresis on a 3130xl Genetic Analyzer (Applied Biosystems), sized against the GeneScan™ 500 LIZ™ Size Standard using the DS-33 dye set, and scored in Geneious.

### Genotype quality filtering and consensus genotype construction

Genotyping errors were minimised using a multiple-tube approach (Taberlet et al., 1996). Consensus genotypes followed the n/2 criterion (Benschop et al., 2013), retaining an allele only when observed in at least half of successful PCR replicates. This criterion was applied together with genotype-specific minimum requirements: heterozygous genotypes required each allele in at least two independent PCRs, whereas homozygous genotypes required the same allele in at least three PCRs (Tumendemberel et al., 2019). Two replicates were initially performed per locus; additional replicates were added when results were inconsistent or failed to meet consensus criteria (Adams & Waits, 2006; Tsaparis et al., 2015). A mean of three replicates per locus was performed (range 2–5), with effort adjusted to DNA quality and replicate consistency.

### Individual identification and genotype validation

Unique individuals were identified from the complete sample-level dataset using Cervus 3.0.7 (Kalinowski et al., 2007). Candidate matches required at least eight matching loci; potential fuzzy matches sharing at least six loci were manually evaluated against replicate-level data to identify scoring errors or allelic dropout. As a cross-method check, samples collected in the Czech Republic and western Slovakia during WY 2023/24 were also analysed in allelematch 2.5.4 (Galpern et al., 2012). The *amUniqueProfile()* function suggested alleleMismatch = 7, but because this estimate generated a cautionary warning, results were interpreted together with Cervus matches and the original PCR replicates.

### Marker discriminatory power

The full multilocus probability of identity among siblings (PID_sib_) was calculated for the complete dataset of 981 unique individuals using the *pid_calc* function in PopGenUtils (Tourvas et al., 2023). To evaluate the effect of marker number, 100 random locus subsets were generated for each subset size from 2 to 20 loci, and multilocus PID_sib_ values were log₁₀-transformed for visualisation. A cumulative analysis was also used to illustrate the decline in PID_sib_ as loci were added to the marker panel.

### Relatedness estimation and dataset pruning

Pairwise relatedness was estimated before population-level analyses using the R package related, with Wang’s estimator as the primary measure (Pew et al., 2015; Wang, 2002). To reduce bias caused by allele-frequency differences in a structured dataset, individuals were assigned to their highest-Q cluster from a preliminary Structure analysis and relatedness was estimated separately within dominant-ancestry groups (Wang, 2011). Individuals showing mixed ancestry were retained and assigned to their highest-Q cluster solely for this purpose. Relatedness was also estimated across the complete dataset to detect close relatives assigned to different ancestry groups. Lynch–Ritland and Queller–Goodnight estimators were calculated as complementary measures (Lynch & Ritland, 1999; Queller & Goodnight, 1989).

Candidate Wang’s r thresholds of 0.45 and 0.55 produced large connected networks that appeared to include broader kin structure. A conservative empirical threshold of r ≥ 0.65 was therefore used to identify the strongest related pairs and networks. One individual was removed from each connected pair or network, prioritising individuals with greater genotyping missingness. The relatedness-pruned dataset was used for subsequent population-genetic analyses.

### Genetic diversity, differentiation and private allelic richness

Observed heterozygosity (H_O_), within-region gene diversity (H_S_) and F_IS_ were calculated for each geographic region using hierfstat (Goudet, 2005). Pairwise Weir–Cockerham F_ST_ estimates and 95% confidence intervals were obtained from 999 bootstrap resamples across loci. Raw private alleles were identified using private_alleles in poppr (Kamvar et al., 2014) and treated descriptively because sample sizes differed among regions. Allelic richness (A_r_) and private allelic richness (PA_r_) were rarefied to 30 gene copies in HP-Rare 1.0 (Kalinowski, 2005), corresponding to the smallest locus-specific number available across regions, and averaged across loci. Rarefied A_r_ was recalculated with allelic.richness in hierfstat as a consistency check.

### Population structure analyses

Population structure was inferred in Structure v2.3.4 (Pritchard et al., 2000) using the relatedness-pruned dataset. Analyses used the admixture model with correlated allele frequencies and no prior population information. K = 1–10 was evaluated with ten replicate runs per K, each comprising 200,000 burn-in iterations followed by 1,000,000 MCMC iterations. Support for alternative K values was assessed in Structure Selector using the Puechmaille estimators, ΔK and mean LnP(K) (Evanno et al., 2005; Li & Liu, 2018; Puechmaille, 2016); the six geographic regions were used as sampling groups for the Puechmaille estimators. Replicates were aligned and summarised in Clumpak (Kopelman et al., 2015).

To describe broad-scale ancestry patterns, we focused on the K = 3 level of population structure. The resulting ancestry components corresponded predominantly to Carpathian, Alpine and Central European populations based on reference membership profiles and geographic distributions. Individuals were classified as showing mixed ancestry when at least two components had Q ≥ 0.20, using a threshold informed by simulation-based assessments of Structure ancestry classification (Vähä & Primmer, 2006); sensitivity was evaluated using maximum individual membership coefficients (Qmax) < 0.75, < 0.80 and < 0.90. For each of the six geographic regions, we summarised the frequency of mixed-ancestry individuals and the distribution of Qmax, including the median and interquartile range. Regional heterogeneity in the frequency of mixed ancestry was evaluated using a conditional Monte Carlo test with fixed marginal totals, followed by pairwise Fisher’s exact tests with Holm correction for multiple comparisons. These operational categories were used to summarise patterns of mixed ancestry and were not interpreted as evidence of specific admixture histories or generations. The complete dataset was analysed under identical settings to assess the effect of relatedness pruning.

DAPC was performed in adegenet as a multivariate approach to assess genetic structure independently of model-based ancestry inference (Jombart, 2008; Jombart et al., 2010). Unsupervised clustering was performed using *find.clusters* and the Bayesian Information Criterion, with 100 principal components retained during clustering; supervised DAPC used the six geographic regions as groups. The number of retained principal components was selected with *xvalDapc* using 50 replicates, 90% training and 10% validation data, and mean assignment success across groups. Data were centred but not scaled, and the maximum available number of discriminant functions was retained. Additional fixed-K analyses for K = 2–10 used 30 cross-validation replicates under otherwise identical settings.

### Individual assignment and recent immigration inference

Individual assignment and exclusion tests were performed in GeneClass2 (Piry et al., 2004) using Alpine, Central European and Carpathian candidate reference populations defined a priori from geographic origin, independently of Structure results and of the non-reference individuals subsequently tested. To balance the reference sets, the Carpathian panel was reduced to a geographically representative subset of Slovak genotypes, prioritising individuals with the fewest missing loci. The initial panel comprised 130 individuals (27 Alpine, 53 Central European and 50 Carpathian) and was evaluated by self-assignment using the Bayesian criterion of Rannala and Mountain (1997), with probabilities estimated by Monte Carlo resampling of 10,000 simulated individuals following Paetkau et al. (2004) and α = 0.01. Individuals whose top-ranked population differed from their geographic origin or that were excluded from their population of origin were removed. This screening was used solely to curate a conservative and internally consistent reference panel and was not treated as an independent validation of assignment accuracy. The final reference panel comprised 123 individuals (26 Alpine, 49 Central European and 48 Carpathian). After the panel was fixed, the remaining 700 non-reference individuals, none of which contributed to reference-panel definition, were evaluated against it using the same settings. Assignment was unique when only one candidate population had a probability ≥ 0.01, ambiguous when multiple populations exceeded this threshold, and unassigned when all three were excluded.

Spatial population structure was investigated separately for nuclear microsatellite and mitochondrial DNA data using the R package geneland (Guillot et al., 2005a,b). All models were spatial, with no additional uncertainty assigned to sampling coordinates. The maximum Poisson–Voronoi process rate was set to the number of analysed individuals, and the maximum number of nuclei was set to three times this value.For the nuclear analysis, an exploratory analysis allowing K to vary from one to ten identified K=7 as the modal solution. We subsequently conducted a final analysis with K fixed at seven using the uncorrelated allele-frequency model. Ten independent MCMC runs of 200,000 iterations were performed, sampling every 100th iteration and thus retaining 2,000 states per run. The first 400 retained states (equivalent to 40,000 iterations; 20%) were discarded as burn-in. Null-allele filtering was disabled. The run with the highest mean post-burn-in log-posterior density was selected for post-processing. For the mitochondrial analysis, haplotypes were supplied as a single haploid locus using the geno.hap argument. An exploratory analysis allowing K to vary from one to ten was performed under the spatial, uncorrelated allele-frequency model. Across the ten exploratory runs, K=7 was the most frequently observed modal value. A final mitochondrial analysis was therefore conducted with K fixed at seven. Ten additional independent runs were performed using the same MCMC, spatial and allele-frequency settings, and the run with the highest mean post-burn-in log-posterior density was selected for post-processing.For both marker systems, mixing of the selected run was assessed by visual inspection of the post-burn-in log-posterior and number-of-nuclei traces, while between-run variation was evaluated by comparing post-burn-in posterior summaries. Selected runs were post-processed on a 300×300-pixel spatial grid.

Recent immigration among the three principal wolf populations was estimated in Ba3Msat v3.0.5 (Wilson & Rannala, 2003) using the relatedness-pruned dataset. The Alpine population comprised geographic region 1, the Central European population pooled the four lowland-related regions 2–5, and the Carpathian population comprised region 6. To minimise effects of unequal sample sizes, we analysed 20 balanced subsets of 27 individuals per population. Each subset was analysed in ten independent chains of 100 million iterations, with 50 million iterations discarded as burn-in and parameters sampled every 5,000 iterations. Convergence was assessed using trace plots, rank-normalised split-R^, and bulk and tail effective sample sizes. Recent immigration was summarised across balanced subsets; full sampling design, diagnostics and sensitivity analyses are provided in Supplementary Methods S2.

### Integration of mitochondrial haplotypes with autosomal ancestry and individual assignment

Mitochondrial haplotypes were linked by genotype identifier to Q coefficients from the relatedness-pruned Structure K = 3 analysis. Of 284 individuals with mtDNA data, 253 were retained in the linked analysis. Haplotypes were grouped descriptively according to their predominant geographic distributions: HW22 as Alpine-associated, HW06 and HW14 as Carpathian-associated, and HW01 and HW02 as Central European-associated. HW03 and HW11 were retained in descriptive haplotype analyses but treated as unclassified for maternal–autosomal concordance because they were not considered sufficiently diagnostic of any of the three focal populations. For the concordance analysis, the predominant autosomal component was defined as the component with the highest individual Q value, and mixed-ancestry classifications were taken from the Structure-based analysis described above.

The linked mtDNA–Structure dataset was combined with GeneClass2 results. Of the 253 individuals, 84 belonged to the reference panel and 169 were non-reference individuals. Concordance involving GeneClass2 was evaluated only among non-reference individuals and, for direct comparisons, only among individuals uniquely assigned at α = 0.01; ambiguous and unassigned individuals were summarised separately. Maternal–autosomal concordance required correspondence between the haplotype-associated background and predominant Structure component, whereas three-way concordance additionally required the same unique GeneClass2 assignment. Individual-level Structure coefficients, mtDNA haplotypes, GeneClass2 results, derived ancestry classifications and regional admixture summaries are provided in Supplementary Data S1.

## Results

### Haplotype distribution

Mitochondrial haplotypes were available for 676 focal and reference individuals (Fig. S2) and showed pronounced spatial structure (Fig. 1a, Fig. 2a). Temporal changes in the spatial occurrence of mitochondrial haplotypes between 2006 and 2026 are shown in Fig. S3. The Central European population is dominated by the haplotype HW01 with a wide occurrence, which, in addition to the Baltic region and the Central European lowland, also dominates the Bohemian Massif and extends into the Central German Uplands and forms enclaves in the Carpathians. Haplotypes HW06 and HW14 dominate in the Carpathians, with HW14 recently spreading into the northern part of the Bohemian Massif. The Alpine population is typical by the occurrence of haplotype HW22, which is currently also spreading to the Central German Uplands and Bohemian Massif.

**Fig. 2.**
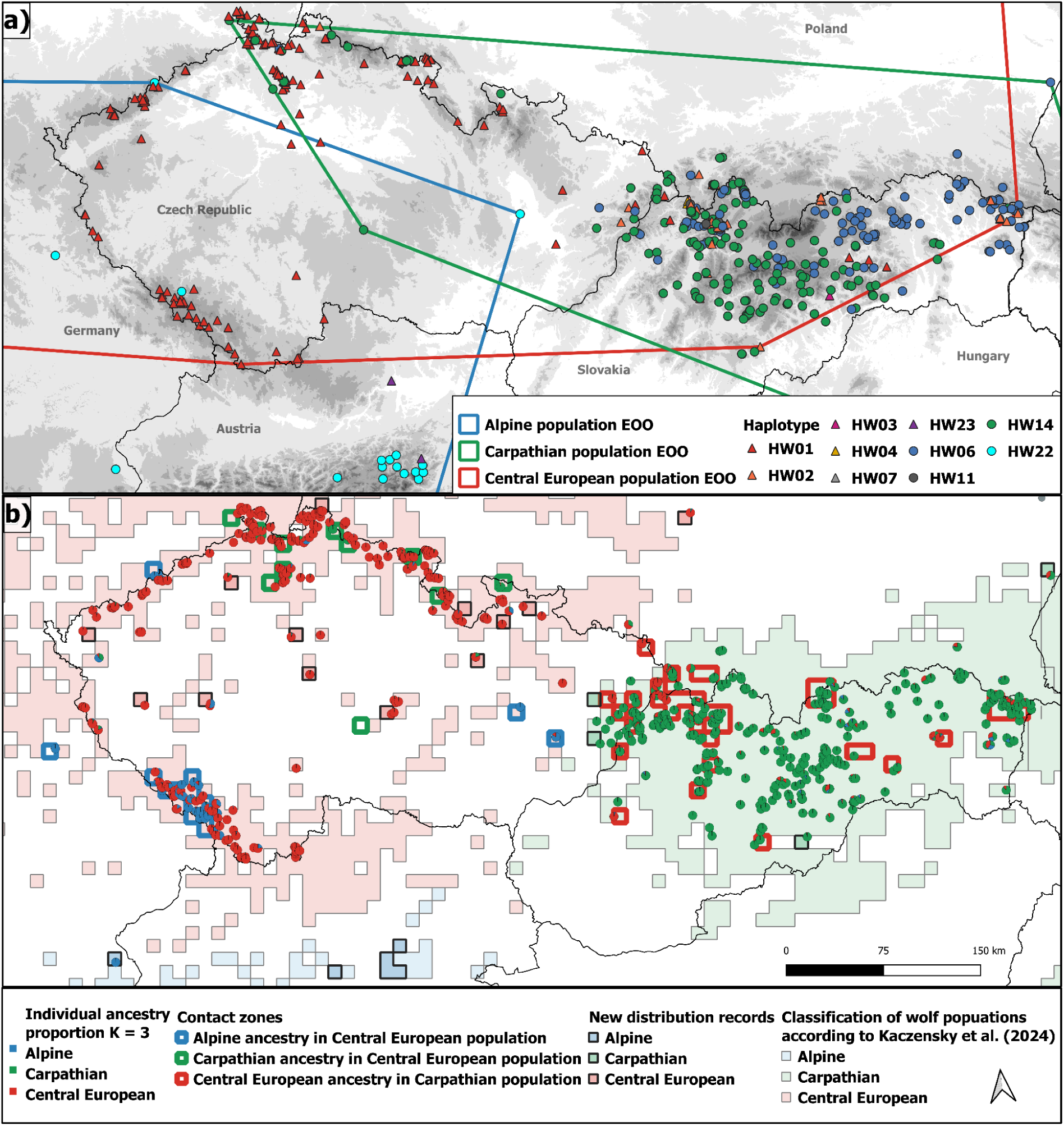
Detailed view of wolf population ranges and genetic structure within the population contact zones in the Czech Republic and Slovakia. **(a)** Individual mtDNA haplotypes are shown by coloured symbols, and maximum spatial extents of the Alpine, Carpathian and Central European populations are represented by minimum convex polygons (MCPs) derived from population-associated mtDNA haplotypes. **(b)** Wolf populations according to Kaczensky et al. (2024) overlaid with individual Structure membership coefficients for K = 3. Each pie chart represents an individual, with sector proportions showing ancestry coefficients inferred from multilocus microsatellite genotypes; colours denote membership in the predominantly Carpathian, Alpine and Central European genetic components.

### Genotyping quality and individual identification

A total of 3,818 biological samples collected in the Czech Republic and western Slovakia yielded 1,903 multilocus wolf genotypes (49.8% genotyping success), representing 524 unique focal individuals. Combined with 457 reference individuals, these formed the complete microsatellite dataset of 981 individuals (517 males, 403 females and 61 of unknown sex). The flow of samples and individuals through the analytical datasets is summarised in Fig. S2. In the WY 2023/24 subset used for cross-method comparison, Cervus combined with replicate-level verification identified 144 unique individuals, whereas allelematch identified 147, a difference of three individuals (approximately 2%).

### Marker discriminatory power

The full 20-locus PIDsib for the complete dataset was 1.98 × 10⁻⁸. In both the cumulative and random-subset analyses, PID_sib_ declined below 0.001 for more than eight loci, indicating sufficient discriminatory power to distinguish closely related individuals (Figs. S4–S5).

### Relatedness pruning

Thresholds of Wang’s r ≥ 0.45 and r ≥ 0.55 produced large connected networks, whereas r ≥ 0.65 identified smaller components representing the strongest relatedness relationships. Pruning reduced the complete dataset from 981 to 823 individuals: 27 from the Alps, 16 from the Central German Uplands, 53 from the Central European lowland, 34 from the Baltic region, 326 from the Bohemian Massif and 367 from the Carpathians.

### Genetic diversity, differentiation and allelic richness

Genetic diversity was highest in the Baltic region (H_O_ = 0.70, H_S_ = 0.73) and lowest in the Alps (H_O_ = 0.58, H_S_ = 0.64). The Carpathians had the highest mean F_IS_ (0.14; Table S2).

Rarefied allelic richness was highest in the Baltic region (A_r_ = 5.88), followed by the Carpathians (A_r_ = 5.53), and lowest in the Alps (A_r_ = 4.39; Table S2). Private allelic richness showed a similar pattern, with the highest values in the Baltic region (PA_r_ = 0.52) and the Carpathians (PA_r_ = 0.43), and the lowest value in the Central German Uplands (PA_r_ = 0.04). Raw private-allele counts, treated descriptively because of unequal sample sizes, were highest in the Carpathians (16 alleles across nine loci), followed by the Bohemian Massif (seven alleles across six loci; Table S2).

Pairwise F_ST_ ranged from 0.004 to 0.176. Differentiation was lowest between the Bohemian Massif and the Central European lowland (F_ST_ = 0.004, 95% CI: 0.001–0.008) and substantially higher between the Bohemian Massif and Carpathians (F_ST_ = 0.121), the Alps and Carpathians (F_ST_ = 0.160), and the Alps and Central European lowland (F_ST_ = 0.176; Table S3).

### Population genetic analyses

Support for the number of Structure clusters differed among estimators. In both the complete and relatedness-pruned datasets, ΔK supported K = 2, whereas mean LnP(K) increased towards K = 10. The Puechmaille estimators supported K = 4 in the complete dataset and K = 3 or K = 4 in the relatedness-pruned dataset, depending on the estimator (Fig. S6). Because the principal large-scale pattern was consistent between datasets, the relatedness-pruned dataset was used for biological interpretation.

Structure revealed hierarchical genetic differentiation. At K = 2, the main split separated the Carpathian component from the remaining samples. At K = 3, the Alpine component separated from the broader non-Carpathian background, yielding three geographically interpretable components corresponding predominantly to the Alpine, Central European and Carpathian populations. The distribution of nuclear genotype enclaves outside the main range mirrored the mitochondrial data (Fig. 1b, Fig. 2b).

Using the criterion Q ≥ 0.20 in at least two components, 67 of 823 individuals (8.1%) showed mixed ancestry. Carpathian–Central European ancestry was most frequent (36 individuals), followed by Alpine–Central European ancestry (24); four individuals combined Alpine and Carpathian components and three exceeded Q ≥ 0.20 in all three components. Mixed ancestry was spatially heterogeneous among geographic regions (conditional Monte Carlo test, p < 10⁻⁶), with a markedly higher frequency in Baltic region (17/34; 50.0%) than elsewhere (0–8.0%). In the Bohemian Massif, 22 of 26 mixed-ancestry individuals combined Alpine and Central European components, whereas 17 of 20 in the Carpathians combined Carpathian and Central European components. The alternative thresholds identified 77 individuals (9.4%) with Qmax < 0.80 and 133 (16.2%) with Qmax < 0.90. The strong regional contrast was retained under alternative thresholds of Qmax < 0.75, < 0.80 and < 0.90 (Supplementary Data S1). At K = 4 and above, additional subdivision occurred mainly in eastern Poland, eastern Slovakia and within the Central European component. Structure plots for K = 1–10 and spatial representations of selected K values are provided in Figs. S7 and S8.

In the unsupervised DAPC workflow, the preceding K-means clustering step based on the Bayesian Information Criterion showed the largest reduction in BIC between K = 1 and K = 2, although BIC continued to decrease at higher K values. We therefore retained K = 2 as the most parsimonious representation of the dominant broad-scale genetic split (Fig. S9a). Cross-validation retained 10 principal components for the subsequent DAPC. The cluster comprising 384 individuals included 351 of 367 individuals from the Carpathians, whereas the cluster comprising 439 individuals contained most individuals from the remaining geographic regions (Fig. S9b). In the supervised DAPC, cross-validation retained 60 principal components. The Carpathians and the Alps were differentiated mainly along the first and second discriminant axes, respectively, whereas the other four regions overlapped substantially (Fig. S10). Fixed-K unsupervised DAPC analyses showed a similar hierarchical pattern. At K = 3, the clusters broadly reflected differentiation among Carpathian, Alpine and Central European affinities, whereas higher K values produced additional, partly overlapping subdivision (Fig. S11).

### Sympatric zones

The current expansion of these populations results in the formation of relatively broad zones where their co-occurrence - and in some cases admixture - takes place, encompassing a broad west-east belt across the study area (Fig. 1), with core areas studied in greater detail in the Bohemian Massif and Western Carpathians (Fig. 3). One zone encompasses the west-east contact line between the Central European lowland and the mountains of the Bohemian Massif and the Carpathians, where the Central European and Carpathian populations intermix (Fig. 3). The second zone, encompassing the contact between the Central European and Alpine populations, has its core in the Bohemian-Bavarian forest.

**Fig. 3.**
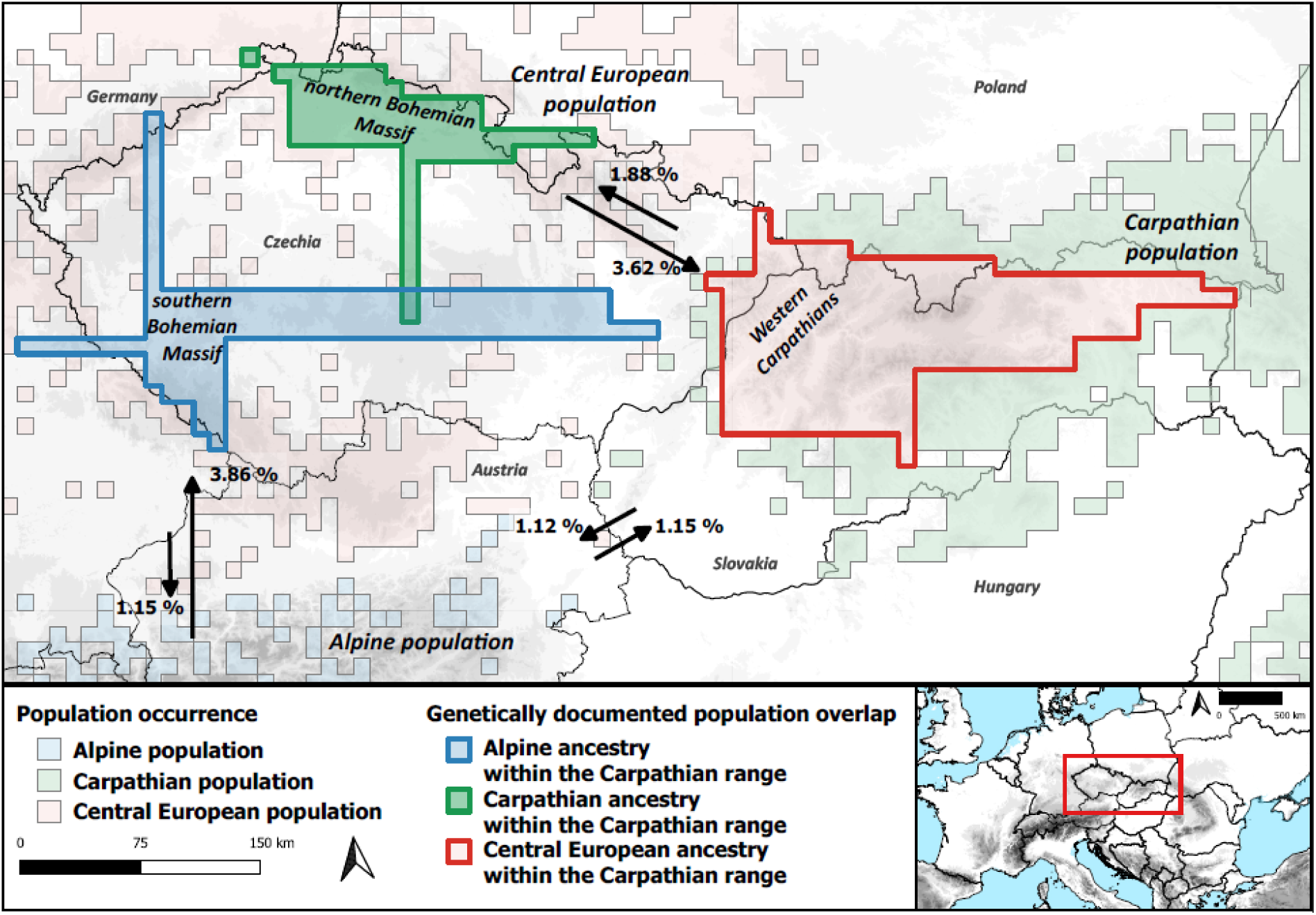
Estimates of maximum extents of interpopulation sympatric zones displayed as orthogonal convex hulls based on occupied EEA grid squares corresponding to population enclaves formed by dispersal outside the range of the source population, based on records of foreign mtDNA haplotypes (representing dispersing animals or backcrosses with mtDNA introgression) or admixed nuclear genotypes with q > 0.75 (corresponding to dispersing animals or first backcross generation). Population distribution is displayed in the EEA grid (Kaczensky et al., 2024 and present study). Gene flow estimates displayed as arrows with respective values.

### Individual assignment and recent immigration inference

Following reference-panel curation, all 123 reference individuals had their geographic population of origin as the top-ranked candidate. Under the more restrictive unique-assignment criterion, 116 individuals (94.3%) were uniquely assigned and seven (5.7%) were ambiguous; none were unassigned. All ambiguous individuals originated from the Central European panel and also showed support for the Carpathian reference population. Among the 700 non-reference individuals, 564 were uniquely assigned at α = 0.01, 38 were ambiguous and 98 were unassigned. Unique assignments comprised 275 Central European, 274 Carpathian and 15 Alpine individuals. GeneClass2 assignments showed a strong geographic pattern (Table S4). In the Bohemian Massif, 255 of 326 non-reference individuals were assigned to the Central European population, whereas 269 of 319 individuals from the Carpathians were assigned to the Carpathian population. The Central German Uplands included ten Alpine and four Central European assignments among 16 individuals. In the Baltic region, 27 of 34 individuals were unassigned, indicating that the three candidate reference populations did not adequately represent the genetic variation in this region.

geneland identified K=7 as the modal solution for both the mitochondrial and microsatellite datasets (Fig. S12). At the broad geographical scale, both marker systems recovered three principal genetic components corresponding to the Alpine, Carpathian and Central European wolf populations. The microsatellite analysis revealed additional fine-scale structure within these broader populations. Wolves from the Bavarian Forest formed a distinct spatial cluster, while the Carpathian population was divided into several geographically structured subclusters. The mitochondrial analysis showed a broadly concordant regional pattern. Overall, the two marker systems recovered similar broad-scale spatial structure but differed in local boundaries and subdivisions. The geographical localization of several mitochondrial clusters is consistent with restricted female-mediated gene flow and the persistence of regional maternal lineages, whereas the microsatellite pattern integrates gene flow mediated by both sexes.

At the population level, Ba3Msat showed strong convergence across all balanced subsets (maximum split-R^ = 1.00012; minimum bulk and tail effective sample sizes = 50,955 and 54,178, respectively). The largest mean recent-immigration estimates were from the Alpine to the Central European population (3.86%; range of subset means 1.83–8.95%) and from the Central European to the Carpathian population (3.62%; 1.29–7.56%; Table S5, Fig. 4). Alpine-to-Central European immigration exceeded the reciprocal estimate in all 20 subsets, whereas Central European-to-Carpathian immigration exceeded the reverse direction in 17 of 20 subsets (Fig. 4). The remaining directional estimates were lower, ranging from 1.12% to 1.88% (Table S5). The principal patterns were retained under alternative treatments of eastern Poland (Supplementary Methods S2).

**Fig. 4.**
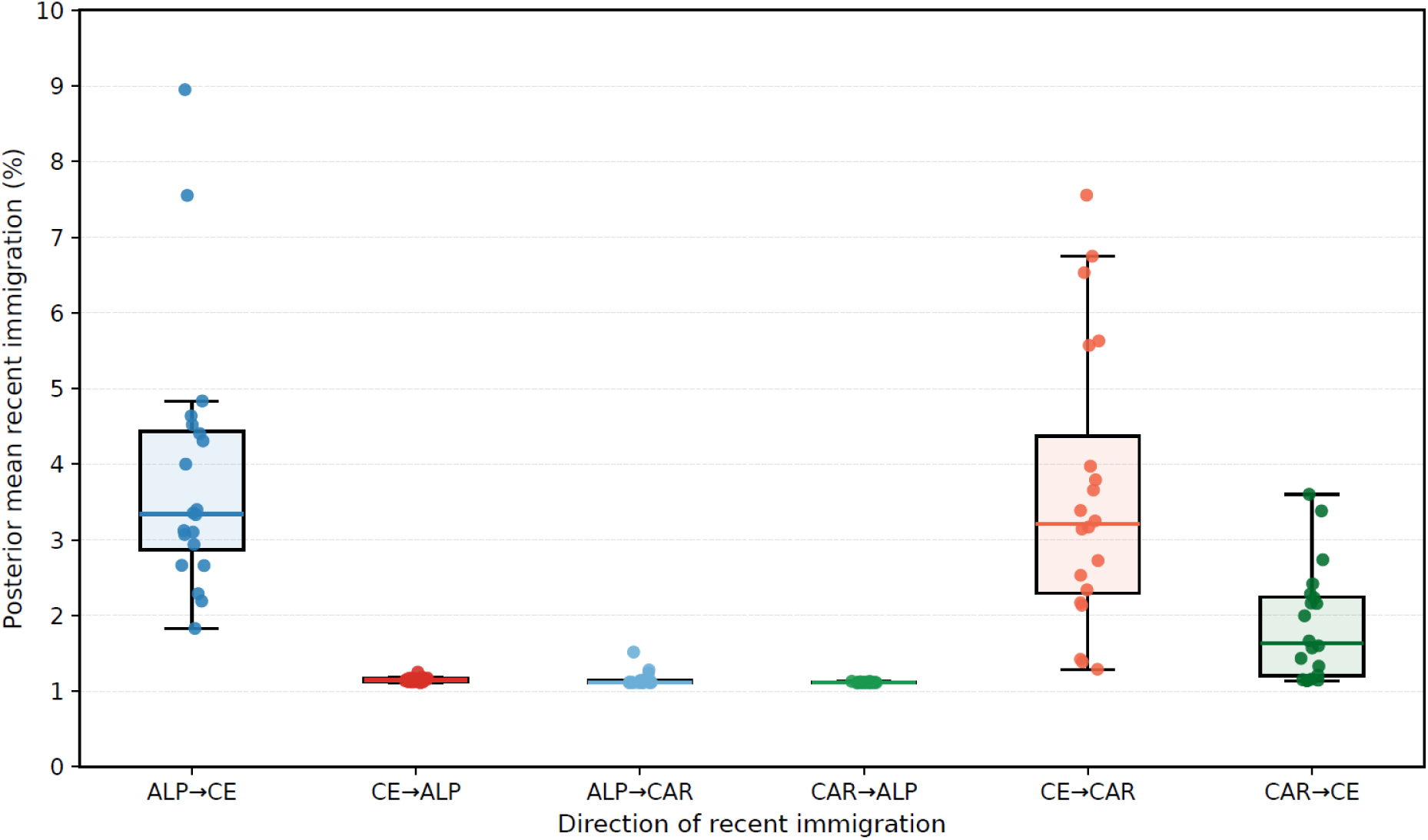
Sensitivity of Ba3Msat recent-immigration estimates to balanced subsampling. Points represent posterior mean recent-immigration proportions for each of 20 balanced subsets comprising 27 individuals per population; boxplots summarise variation among subset-specific posterior means. Arrows indicate source-to-recipient direction. Immigration from the Alpine (ALP) to the Central European population (CE) exceeded the reciprocal direction in all 20 subsets, whereas Central European-to-Carpathian (CAR) immigration exceeded the reverse direction in 17 of 20 subsets.Sensitivity of Ba3Msat recent-immigration estimates to balanced subsampling. Points represent posterior mean recent-immigration proportions for each of 20 balanced subsets comprising 27 individuals per population; boxplots summarise variation among subset-specific posterior means. Arrows indicate source-to-recipient direction. Immigration from the Alpine (ALP) to the Central European population (LOW) exceeded the reciprocal direction in all 20 subsets, whereas Central European-to-Carpathian (CAR) immigration exceeded the reverse direction in 17 of 20 subsets.

### Concordance among mitochondrial haplotypes, Structure ancestry and GeneClass2 assignment

Mitochondrial haplotypes were linked to Structure K = 3 membership coefficients for 253 of 284 individuals with mtDNA data (89.1%); the remaining 31 had been removed during relatedness pruning. After excluding individuals carrying the unclassified HW03 (n = 2) and HW11 (n = 11) haplotypes from maternal-background comparisons, 219 of 240 individuals (91.3%) showed correspondence between their haplotype-associated maternal background and predominant Structure component. All 40 HW22 individuals showed predominant Alpine membership, whereas HW14 and HW06 were predominantly associated with the Carpathian component and HW01 and HW02 with the Central European component. HW11 nevertheless showed a strong autosomal association with the Central European component: all 11 carriers had predominant Central European Structure membership (Table S6). Nineteen of the 253 linked individuals (7.5%) showed mixed Structure ancestry. In 17 cases, the component corresponding to the maternal background remained among those with Q ≥ 0.20. Maternal and predominant autosomal backgrounds differed in 21 of 240 individuals (8.8%): 11 combined Central European-associated mtDNA with Alpine membership, seven combined Carpathian-associated mtDNA with Central European membership, and three showed the reverse Central European–Carpathian combination. Ten of the 11 individuals with Central European-associated mtDNA and Alpine membership occurred in or near the

Bohemian–Bavarian Forest contact area. Among 169 linked non-reference individuals, 127 were uniquely assigned in GeneClass2, eight were ambiguous and 34 were unassigned. Among uniquely assigned individuals, GeneClass2 corresponded to the predominant Structure component in 125 cases (98.4%). Among individuals with a classified maternal background, GeneClass2 corresponded to the maternal background in 121 of 126 cases (96.0%), and all three classifications agreed in 120 of 126 cases (95.2%). Of 18 non-reference individuals showing mixed Structure ancestry, 11 were unassigned by GeneClass2, one was ambiguous and six were uniquely assigned. Complete individual-level concordance results and alternative Q-threshold classifications are provided in Supplementary Data S1.

## Discussion

Range expansion during population recovery can substantially reshape genetic structure (Excoffier et al., 2009), particularly when historically differentiated populations come into secondary contact (Hewitt, 2000). In such systems, an important question is whether historical population structure persists despite renewed connectivity, or becomes progressively eroded as dispersal and admixture increase. The ongoing recovery of wolves in Central Europe provides an opportunity to examine this process across the contact of several differentiated population backgrounds. Here, we evaluated how these backgrounds are spatially distributed, to what extent they remain genetically distinguishable, and how contemporary contact is reflected in mixed ancestry and recent immigration. Overall, our results point to a spatially structured system of secondary contact in which historical differentiation persists despite increasing population overlap.

The three principal genetic backgrounds were consistently recovered by both model-based and multivariate analyses, in agreement with previously documented mitochondrial and nuclear differentiation among European wolf populations (Czarnomska et al., 2013; Hulva et al., 2018, 2024; Pilot et al., 2010; Stronen et al., 2013). Population structure was hierarchical, and additional subdivision emerged at higher K values, partly in correlation of isolation of recognized geographical regions (Fig. 1b). Although rarefaction reduces the effect of unequal sample size (Kalinowski, 2004, 2005; Leberg, 2002), regional richness estimates remain sensitive to spatial coverage. Uneven sampling and isolation by distance can affect Bayesian clustering (Frantz et al., 2009; Puechmaille, 2016), and ancestry components do not necessarily represent discrete biological populations or literal historical sources (Lawson et al., 2018). The threshold-based Q classification should likewise be regarded as a descriptive measure of intermediate ancestry rather than as evidence of discrete admixture categories or a particular hybrid generation. Broad agreement with DAPC nevertheless supports using the three principal backgrounds as a framework for describing contemporary population structure and secondary contacts.

### Population-specific patterns during recolonisation

Elevated diversity within Baltic region may reflect regional substructure and contributions from eastern or Baltic-related genetic backgrounds, in agreement with the complex population structure previously described in Polish and eastern European wolves (Czarnomska et al., 2013; Stronen et al., 2013; Szewczyk et al., 2019) and denotes the area as source, from which the Central European population expanded westward (Jarausch et al., 2021; Szewczyk et al., 2019). This may also reflect larger effective population size, historical or recent connection to larger and more diverse populations in Asia (not represented in the present dataset), gene flow from Carpathians, finer-scale regional structure and other factors. The particularly high proportion of unassigned individuals in eastern Poland may reflect mixed ancestry or genetic variation not represented by the three reference populations, because assignment and exclusion probabilities depend on the differentiation, composition and sampling of candidate populations (Cornuet et al., 1999; Manel et al., 2005; Paetkau et al., 2004). For example, the occurrence of the HW3 haplotype in the south of the area can be attributed to gene flow from the Pontic (Ukrainian Steppe) population (Hulva et al., 2018; Szewczyk et al., 2021). Broader reference sampling would be required to distinguish among these alternatives.

Within the Central European population, DAPC showed substantial overlap between the Bohemian Massif and Central European lowland, consistent with broad genetic continuity across the recently recolonised range. This pattern, together with moderate allelic richness and comparatively little private variation in the Bohemian Massif, is consistent with recent westward expansion from western Poland through Germany and adjacent regions (Jarausch et al., 2021; Szewczyk et al., 2019), as well as with previous evidence that the Central European population has expanded across both lowland and mountainous landscapes (Szewczyk et al., 2019, 2021).

Rarefied allelic and private allelic richness were high in the Carpathians, in agreement with previous evidence of comparatively high genetic diversity in eastern and Carpathian wolf populations (Jan et al., 2023; Stronen et al., 2013). The comparatively high F_IS_ in the Carpathians may likewise partly reflect geographic substructure and a Wahlund effect rather than inbreeding alone (Garnier-Géré & Chikhi, 2013). Finer-scale regional structure was also detected in eastern Slovakia, with broadly concordant patterns inferred by Structure and geneland. The subdivision in eastern Slovakia resembles the east–west genetic discontinuity reported in Carpathian brown bears, providing a parallel example of regional genetic structure in another Carpathian large carnivore (Straka et al., 2012).

In contrast, the Alpine population had the lowest rarefied allelic richness, consistent with the documented founder-driven expansion of the refugial Italian population from the Apennines (Fabbri et al., 2007).

### Secondary zones of sympatry and gene flow

Observed patterns of sympatry are spatially mosaic and may partly reflect typical wolf spatial ecology with long-distance dispersal by subadults (Kojola et al., 2006; Morales-González et al., 2022). Exact proportions of “pure” individuals from parental populations and their hybrids of particular generation and backcrosses vary among regions. Temporal changes in these mosaic patterns (Fig. S3) may reflect local demographic turnover or the gradual erosion of enclave-specific ancestry signals through introgression and/or mortality (e.g. in Carpathians, whole packs are often culled). However, further data are necessary to reduce temporal variation in sampling and detection in some regions.

Among uniquely assigned non-reference individuals, GeneClass2 assignments corresponded closely to predominant ancestry inferred by structure, although this agreement should not be regarded as independent validation because both analyses used the same microsatellite loci. Broad concordance between mitochondrial haplotypes and autosomal ancestry further supports the geographic interpretation of the three principal genetic backgrounds, whereas maternal–autosomal discordance was concentrated around population interfaces and provides complementary evidence of secondary contact. Central European-associated mtDNA combined with predominant Alpine ancestry occurred mainly in or near the Bohemian–Bavarian Forest, while discordant Central European–Carpathian combinations occurred around the eastern contact interface. Such discordance is compatible with dispersal followed by introgression, although maternally inherited mtDNA cannot establish the sex, direction or generation of the dispersing ancestors.

At the population level, Ba3Msat estimates indicated asymmetric recent immigration among the three major genetic backgrounds. Interactions between Central European and Carpathian populations were most intense, leading to the formation of two enclaves or satellite populations, comprising Central European wolves in several areas of the Eastern Carpathians and Carpathian wolves in the Sudetes mountains in the northern part of the Bohemian Massif (Fig. 3). These new records substantially extend earlier evidence of dispersal and admixture between these populations (Czarnomska et al., 2013; Hulva et al., 2018; Szewczyk et al., 2019). The occurrence of ancestry components beyond their principal ranges is consistent with dispersal across major landscape features, including the Outer Carpathian depressions with substantial landscape resistance to movements of large mammals including natural (rivers) and anthropogenic barriers (linear infrastructure, settlements). The relationship is asymmetrical with Central European-to-Carpathian immigration exceeding the reverse direction in 17 of 20 subsets (Fig. 4). Importantly, the principal patterns were stable under alternative treatments of eastern Poland. These estimates represent recent immigration proportions among sampled wolves rather than annual migration rates or direct measures of successful reproductive gene flow.

Dispersal between Central European and Alpine populations led to the formation of one enclave or satellite population, comprising the southern Bohemian Massif, particularly the Bohemian–Bavarian Forest. This suggests the permeability of lowland barriers like the Danube valley for the Alpine wolves. Admixture between the Central European and Alpine populations was previously reported (Hulva et al., 2024) and is reflected in a distinct cluster recovered by both Structure and geneland in the present study. Most mixed-ancestry individuals in the Bohemian Massif combined Alpine and Central European components. The Bohemian–Bavarian Forest may act as a stepping stone for further spread of Alpine genetic variation (Fig. S12), although continued monitoring is needed to confirm reproduction and expansion beyond the contact zone. Vagrant animals from the Alps were identified in Central German Uplands and Bohemian Massif. Continuing expansion and long-distance dispersal of Alpine wolves (Marucco et al., 2022, 2023) indicate potential for further occurrence of admixture within contact zone EOO. Immigration from the Alpine into the Central European population was the most consistent directional signal, exceeding the reciprocal estimate in all 20 balanced subsets. Together with the geographic concentration of Alpine ancestry and assignments in the Bohemian–Bavarian Forest, this supports continuing contact across the western population interface, although the population-level model does not localise individual migration events to this region.

The level of gene flow between the Carpathian and Alpine populations was low in both directions, likely due to the geographical configuration and extensive lowland barriers - such as the Pannonian Basin - separating the two populations. However, additional data - including other countries such as Austria - will be needed to more precisely quantify the dynamics of gene flow between these populations.

Despite dispersal capacity, the persistence of spatial associations between genetic backgrounds and geographic regions suggests that behavioral ecology may contribute to maintaining population structure. Ecologically associated genetic differentiation in eastern European wolves has previously been interpreted in terms of natal habitat-biased dispersal (Pilot et al., 2006), and a direct test in Scandinavian wolves found preferential settlement in natal-like habitats among shorter-distance dispersers, although this pattern weakened with increasing dispersal distance (Sanz-Pérez et al., 2018). At the same time, recent demography history and anthropogenic barriers may also have shaped population structure during the expansion of the Central European population (Szewczyk et al., 2019). Within the Bohemian Massif, the availability of transitional habitats may facilitate settlement by wolves from different genetic backgrounds, whereas the region’s relative topographic isolation may contribute to the persistence of the spatially structured ancestry mosaic observed here. These explanations are not mutually exclusive; distinguishing the relative contribution of natal habitat-biased settlement, demographic history and anthropogenic factors and landscape configuration would require individual dispersal histories combined with quantitative environmental data.

### Conservation implications

Whether continued expansion will ultimately lead to progressive genetic homogenisation or to the persistence of a spatially structured admixture mosaic remains unresolved, as well as potential consequences for population fitness (Schumer & Rieseberg, 2026). Current findings based on analyses of a large number of genomes indicate low effective population sizes for European wolves, thereby highlighting their evolutionary vulnerability (Todd et al., 2026). From the perspective of conservation genetics/genomics, an assessment of increasing inter-population admixture could therefore be positive, as it might lead to a heterosis effect. Here, however, we encounter the limits of our knowledge, as we lack information regarding the adaptations of individual populations to differing environmental conditions - such as those observed in North American wolves (Schweizer et al., 2016) - that could potentially lead to outbreeding depression. Continued temporal and transboundary genetic monitoring, extending to other countries within the interpopulation contact zones (e.g., Poland, Germany, and Austria), will be important given the high dispersal capacity of wolves and the potential for these zones to expand rapidly. Integrating genomic and phenotypic data will be important for providing insights into the development of these emerging contact zones and the extent to which admixture may influence the future genetic structure, viability, and conservation management of Central European wolf populations.

The magnitude and causes of anthropogenic wolf mortality can vary among regions and include traffic collisions, legal culling, poaching etc. (Morales-González et al., 2026; Nowak et al., 2021). Management regimes likewise differ among jurisdictions. In Slovakia, a public wolf-hunting scheme based on annual quotas operated during 2014–2019 (Kutal et al., 2024), and quota-based hunting was reinstated in 2025 (Kutal et al., 2025). From a management perspective, the asymmetric recent immigration from the Central European to the Carpathian population raises the possibility of source–sink-like dynamics, in which immigration may supplement populations experiencing higher local mortality (Pulliam, 1988). However, explicit demographic data are needed for testing this hypothesis.

Because the identified areas of sympatry do not coincide with administrative boundaries, individual monitoring units may encompass multiple genetic backgrounds, mixed-ancestry individuals and recent dispersers, while each population may extend across several jurisdictions. These transboundary areas of sympatry call for coordinated genetic monitoring. Harmonised markers, reference datasets and analytical standards are needed to compare genotypes generated by different laboratories and countries, particularly where Central European, Carpathian and Alpine backgrounds occur in close proximity (de Groot et al., 2016; Hindrikson et al., 2017). Transnational monitoring can resolve population processes that cross administrative boundaries, and its feasibility has been demonstrated for large carnivore populations, including wolf monitoring coordinated across eight Central European countries by the Central European Wolf (CEwolf) Consortium and across seven Alpine countries by the Wolf Alpine Group (WAG).

## Acknowledgements

We thank the Nature Conservation Agency of the Czech Republic, particularly Petr Kafka, Jan Lukavský, Václav Tomášek, Jakub Čejka, Pavel Jaška, Petr Kuna, Lenka Lánská and Hana Bednářová; national parks in the Czech Republic; Friends of the Earth Czech Republic - Carnivore Conservation Programme, particularly Michal Bojda, Barbora Černá, Kristýna Chroboková, Jan Drapák, Rostislav Dvořák, Michal Feller, Zuzana Fišerová, Šárka Frýbová, Štěpánka Kadlecová, Jan Koranda, Josefa Krausová, Radek Kříček, Jiří Labuda, Jakub Lalouček, František Moupic, Martin Váňa and the volunteers of the Carnivore Tracking Project; OWAD & OWADIS & REDEMA team (namely Lukáš Žák, Jan Horníček) and WoBoFe team (namely Oldřich Vojtěch sen., Pavla Jůnková Vymyslická); LECA team and the State Nature Conservancy of the Slovak Republic, particularly Mária Apfelová, Jozef Štofík, Peter Drengubiak, Tomáš Flajs, Tibor Pšenák, Radovan Reťkovský and János Csík, for field sampling. We thank CEwolf consortium, within which we carried out the harmonization of genetic markers, enabling transboundary monitoring. We thank Alžběta Báčová, Magdalena Bélová, Markéta Benešová, Klára Demjanovičová, Milena Jindřichová, Monika Ladányiová and Vendula Woznicová for contribution during the labwork. We are also grateful to the Association for Nature “Wolf” for supporting sample collection in Poland.

## Ethics Statement

Ethical review and approval were not required for this study because no live animals were captured, handled, or subjected to experimental procedures for the purposes of the research. Genetic material was obtained from non-invasive samples collected during routine wolf monitoring, swabs taken from livestock carcasses, and tissue samples from wolf mortalities unrelated to the study.

## Data Availability Statement

The processed data supporting the findings of this study are provided in the article and its Supporting Information. The underlying raw data are available from the corresponding author upon reasonable request.

